# Tumour purity as a prognostic factor in colon cancer

**DOI:** 10.1101/263723

**Authors:** Yihao Mao, Qingyang Feng, Peng Zheng, Liangliang Yang, Tianyu Liu, Yuqiu Xu, Dexiang Zhu, Wenju Chang, Meiling Ji, Yongjiu Tu, Li Ren, Ye Wei, Guodong He, Jianmin Xu

## Abstract

Tumour purity is defined as the proportion of cancer cells in the tumour tissue. The impact of tumour purity on colon cancer (CC) prognosis, genetic profile and microenvironment has not been thoroughly accessed. Therefore, clinical and transcriptomic data from three public datasets, GSE17536/17537, GSE39582, and TCGA were retrospectively collected (n = 1248). Tumour purity of each sample was inferred by a computational method based on transcriptomic data. Stage III and MMR-deficient (dMMR) CC patients showed a significantly lower tumour purity. Low purity CC conferred worse survival and tumour purity was identified as an independent prognostic factor. Moreover, high tumour purity CC patients benefited more from adjuvant chemotherapy. Subsequent genomic analysis found that the mutation burden was negatively associated with tumour purity with only APC and KRAS significantly more mutated in high purity CC. However, no somatic copy number alteration event was correlated with tumour purity. Furthermore, immune-related pathways and immunotherapy-associated markers (PD-1, PD-L1, CTLA-4, LAG-3, and TIM-3) were highly enriched in low purity samples. Notably, the relative proportion of M2 macrophages and neutrophils, which indicated worse survival in CC, was negatively associated with tumour purity. Therefore, tumour purity exhibited potential value for CC prognostic stratification as well as adjuvant chemotherapy benefit prediction. The relative worse survival in low purity CC may attribute to higher mutation frequency in key pathways and purity related microenvironmental changing.

**Summary:** Low purity colon cancer patients conferred worse survival and benefited less from adjuvant chemotherapy. The mutation burden was negatively associated with tumour purity. Low purity samples exhibited intense immune phenotype with more M2 macrophages and neutrophils infiltration.

## Introduction

Colorectal cancer (CRC) was estimated to be the third most commonly diagnosed cancer in the USA for both men and women in 2018 and nearly 70% were colon cancer (CC) cases (1). Increasing evidence suggested tumour microenvironment (TME), a collection of cancer cells and neighbouring tumour-associated non-cancerous cells plays a pivotal role in tumour biology (2),(3). Numerous researches supported that tumour-associated stroma takes part in tumour progression, metastasis and response to chemotherapy (4).

Tumour purity is the percentage of tumour cells in TME. Previous studies investigated the relationship between the proportion of cancer cells and prognosis. Using TCGA data, a pan-cancer analysis of tumour purity found high tumour purity predicted better survival in kidney renal clear cell carcinoma and lower-grade glioma (5). In colon cancer (CC), several studies proposed stroma percentage conferred worse survival (6–10). However, previous studies used visual or computer-assisted estimation of tumour and stroma proportion based on hematoxylin and eosin-stained slides, which may introduce human error and subjective bias. In addition, current understanding of the purity-related genomic alterations and microenvironmental changing was still limited.

In our study, we inferred tumour purity of CC patients in public datasets by a computational method based on transcriptomic data. The relationship between tumour purity, clinicopathological characteristics, and prognosis in CC was further investigated. Moreover, gene mutations, somatic copy number alterations (SCNAs), biological pathways, and immune cells infiltration associated with tumour purity in CC were thoroughly explored, which may provide a deeper understanding and help clinical management of CC.

## Materials and methods

### Study population

Public datasets selection criterion was as follows: (1) transcriptomic data (microarray data or RNA Seq data) were available; (2) the basic clinicopathological information (stage and survival information) was available; (3) the sample size was larger than 100. Therefore, three datasets were included in our research (GSE17536/17537, GSE39582, TCGA) (11–14). Furthermore, patients with no matched transcriptomic data or at stage 0 were excluded. Finally, 1248 patients were enrolled in our study (Figure S1). All patients were pathologically diagnosed with primary CC and staged according to the American Joint Committee on Cancer (AJCC) staging system. The clinicopathological information was collected from the corresponding data portal: Gene Expression Omnibus (GEO) repository (https://www.ncbi.nlm.nih.gov/gds/) and Genomic Data Commons (GDC) Data Portal (https://portal.gdc.cancer.gov/). The DNA mismatch repair (MMR) status was determined by immunohistochemistry (IHC) staining. Detailed clinicopathological characteristics of enrolled patients from three datasets were described in Table S1.

### Tumour purity estimation

The MINiML formatted family files containing metadata and Affymetrix HG133plus2 microarray data of GSE17536/17537 and GSE39582 were downloaded from GEO repository. TCGA level 3 RNA-Seq Version 2 RSEM data were obtained from GDC Data Portal. ESTIMATE was a widely used R library for tumour purity prediction (15,16). The expression profile of 141 stroma-related genes and 141 immune-related genes was analysed and tumour purity was estimated by combining stromal and immune scores. By running ESTIMATE on TCGA RNA-Seq and Affymetrix HG133plus2 microarray data, tumour purity of each CC sample can be estimated as described before (16). The result was averaged if a sample had more than one matched transcriptomic profile. The RNA-Seq data of 48 CC cell lines were obtained from Cancer Cell Line Encyclopedia (CCLE, http://www.broadinstitute.org/ccle) for validation of ESTIMATE algorithm (17).

### Genomic analysis

TCGA level 3 RNA-Seq Version 2 RSEM data, segmented somatic copy number alteration (SCNA) data (minus germline CNV), and TCGA level 3 mutation data of version 2016_01_28 were downloaded from GDC Data Portal. Mutation data were analysed and summarized using maftools (18). Differential mutations between groups were accessed applying Fisher’s exact tests. KEGG pathway analyses were performed using DAVID 6.8 for differential mutated genes (19, 20). SCNA events were detected by Genomic Identification of Significant Targets in Cancer (GISTIC) 2.0 using the segmented Affymetrix SNP 6.0 microarray data (21). SCNAs between groups were compared by Fisher’s exact tests. An R package, edgeR, was utilized to perform differential expression analysis (22). Gene set enrichment analysis (GSEA) was performed by the GSEA desktop application v.3.0 using Molecular Signatures Database (MSigDB) v6.1 with 1000 permutations (23, 24). The estimation of immune cell proportions for each sample was performed by CIBERSORT algorithm on TCGA RNA-Seq data or Affymetrix HG133plus2 microarray data using LM22 as a reference expression signature with 100 permutations.

### Statistical Analysis

All statistical analyses were performed using SPSS 22.0 (SPSS Inc., Chicago, IL) and R software, version 3.4.3 (The R Foundation for Statistical Computing, http://www.r-project.org/). Variables between groups were compared by the Student t-tests or one-way analyses of variance (ANOVA) with post hoc pairwise Bonferroni tests. Correlations between continuous variables were evaluated by Spearman correlation analyses. Kaplan-Meier analyses and Log-rank tests were used to evaluate the relationship between groups and overall survival (OS). Univariate and multivariate Cox regression analyses were performed to identify independent prognostic factors, factors with P < 0.1 in univariate Cox regression analyses were further evaluated in the multivariate Cox regression models. A two-sided P < 0.05 was regarded as statistically significant.

### Ethics statement

GSE17536/17537, GSE39582, and TCGA are pubic datasets. Therefore, ethics committee approval was not needed. Neither patient informed consent nor permission to use this data was required to perform the current analyses.

## Results

### Tumour purity and clinicopathological features

Tumour purity of each CC sample was calculated by ESTIMATE algorithm. Among 1248 patients, tumour purity ranged from 0.195 to 0.984 with the median purity of 0.765. The distribution of purity was presented in Figure 1A. 48 CC cell lines were used as references. The purity of CC cell lines ranged from 0.989 to 1.000 with the median purity of 0.998 (Fig. 1B), which validated the robustness of ESTIMATE algorithm. Baseline clinicopathological characteristics of three datasets were shown in Table S1. Furthermore, the correlation between purity and clinicopathological features was evaluated in Figure 1B and Table S2. Tumour purity was significantly associated with TNM stage and MMR status (both *P* < 0.05). Pairwise Bonferroni tests indicated that tumour purity of stage III patients was lower than stage I ones (P = 0.005). MMR-deficient (dMMR) CC patients showed a significantly lower tumour purity compared with MMR-proficient (pMMR) patients (P = 0.009), which was consistent with previous report (8).

**Fig. 1.**
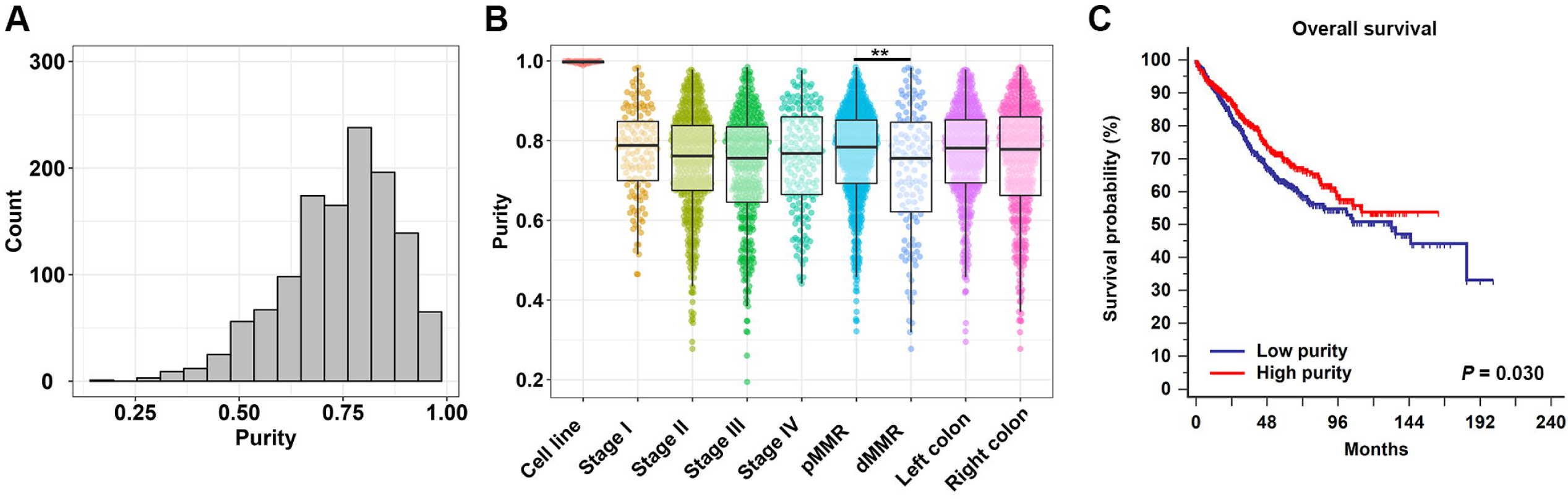
(**A**) The distribution of tumour purity in colon cancer. (**B**) The distribution of tumour purity among TNM stages, MMR status, and primary locations. (**C**) Kaplan–Meier analysis of overall survival showed low purity colon cancer (separated by median purity) conferred worse prognosis. TNM, tumour node metastasis; MMR, DNA mismatch repair; dMMR, MMR-deficient; pMMR, MMR-proficient; **P = 0.009.

### Lower purity conferred worse prognosis in colon cancer

CC Samples were divided into low and high purity groups by median purity value. As accessed by Kaplan–Meier survival analyses and Log-rank tests, the high purity group conferred prognostic benefit (P = 0.030) (Fig. 1C). In further univariate Cox regression analyses, age, TNM stage and tumour purity (as a continuous variable) were associated with OS (all P < 0.05). In multivariate Cox regression analysis, tumour purity was identified as an independent prognostic indicator of OS (P = 0.022, HR = 0.389, 95% CI = 0.174 – 0.870) (Table 1).

**Table 1.** Univariate and multivariate Cox regression analyses for OS in combined dataset

|  | Univariate analysis |  | Multivariate analysis |  |
| --- | --- | --- | --- | --- |
| Factors | HR (95% CI) | *P* | HR (95% CI) | *P* |
| Age (years) |  |  |  |  |
| Increasing years | 1.018 (1.009 - 1.027) | **<0.001** | 1.027 (1.018 - 1.036) | **<0.001** |
| Gender |  |  |  |  |
| Male | 1 (reference) |  |  |  |
| Female | 0.845 (0.683 - 1.046) | 0.122 |  |  |
| Primary site |  |  |  |  |
| Right-sided colon | 1 (reference) |  |  |  |
| Left-sided colon | 0.895 (0.696 - 1.150) | 0.386 |  |  |
| TNM Stage |  |  |  |  |
| I | 1 (reference) |  | 1 (reference) |  |
| II | 1.809 (1.019 – 3.212) | **0.043** | 1.786 (1.005 - 3.174) | **0.048** |
| III | 2.368 (1.333 – 4.206) | **0.003** | 2.502 (1.404 - 4.459) | **0.002** |
| IV | 9.940 (5.578 – 17.712) | **<0.001** | 11.089 (6.207 - 19.811) | **<0.001** |
| MMR status |  |  |  |  |
| MMR-proficient | 1 (reference) |  | 1 (reference) |  |
| MMR-deficient | 0.614 (0.382 – 0.984) | **0.043** | 0.701 (0.435 – 1.128) | 0.143 |
| Purity |  |  |  |  |
| Increasing purity | 0.487 (0.225 – 1.055) | 0.068 | 0.389 (0.174 – 0.870) | **0.022** |
| Chemotherapy |  |  |  |  |
| No | 1 (reference) |  |  |  |
| Yes | 1.015 (0.774 - 1.331) | 0.912 |  |  |

Abbreviations: TNM, tumor node metastasis; OS, overall survival; MMR, DNA mismatch repair; HR, harzard ratio; CI: confidence interval

Subgroup analyses revealed that low tumour purity indicated impaired survival in male (P = 0.010) and pMMR CC patients (P = 0.041) (Figure 2). Interestingly, patients who underwent adjuvant chemotherapy with high tumour purity had significant survival benefit compared with low tumour purity patients (P = 0.016, HR = 0.166, 95% CI = 0.039 – 0.714). Subsequent multivariate Cox regression analyses adjusted for TNM stage identified increasing purity as an independent predictor for chemotherapy benefit (P = 0.005, HR = 0.129, 95% CI = 0.031 – 0.537) (Table S4).

**Fig. 2.**
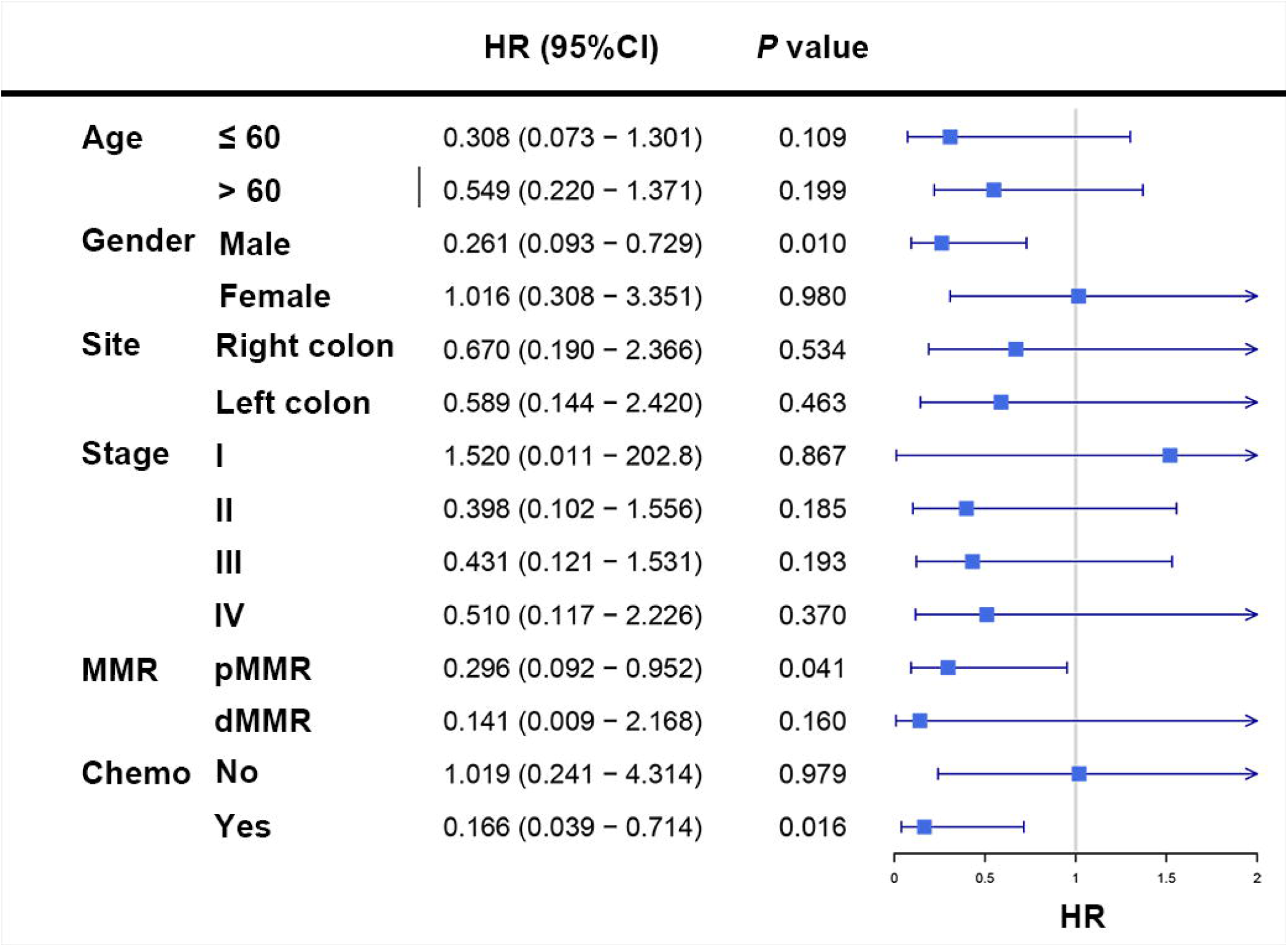
Subgroup analyses revealed that low tumour purity indicated impaired survival in male, pMMR CC patients, and patients with adjuvant chemotherapy. MMR, DNA mismatch repair; dMMR, MMR-deficient; pMMR, MMR-proficient; Chemo: adjuvant chemotherapy; HR: hazard ratio; CI: confidence interval.

### Tumour purity and genomic profile

To unveil the possible mechanisms affecting tumour purity, genomic data including mutation profile and somatic copy number alteration (SCNA) data of TCGA dataset were further analysed. The relationship between mutation profile and purity was evaluated in 394 TCGA patients with available somatic mutation data. Parallel analyses were conducted between three kinds of subgroups (1st vs. 2nd half, 1st vs. 3rd tertile, 1st vs. 4th quarter) respectively. Linear regression analysis showed that tumour purity was negatively correlated with somatic mutations (R = −0.206, P < 0.001) (Figure S2). More mutations were detected in low purity samples (Mean mutation number: 1st vs. 2nd half: 577.0 vs. 269.4; 1st vs. 3rd tertile: 596.5 vs. 217.7; 1st vs. 4th quarter: 707.0 vs. 178.2).

The summary of overall mutation profile of TCGA dataset was illustrated in Figure S3. APC, TP53, TTN and KRAS ranked as top mutated genes as described before (14). The most frequently mutated genes among low and high purity groups were presented in Figure 3A, Figure S4A, and Figure S5A. Most genes including *DNAH7, KMT2D, RYR3, BRAF, IGF2 R*, etc. were found significantly more mutated in low purity group (all P < 0.05). Notably, only *APC* and *KRAS* mutations were more frequently detected in high purity group (both P < 0.05) (Figure 3B, Figure S4B, and Figure S5B).

**Fig. 3.**
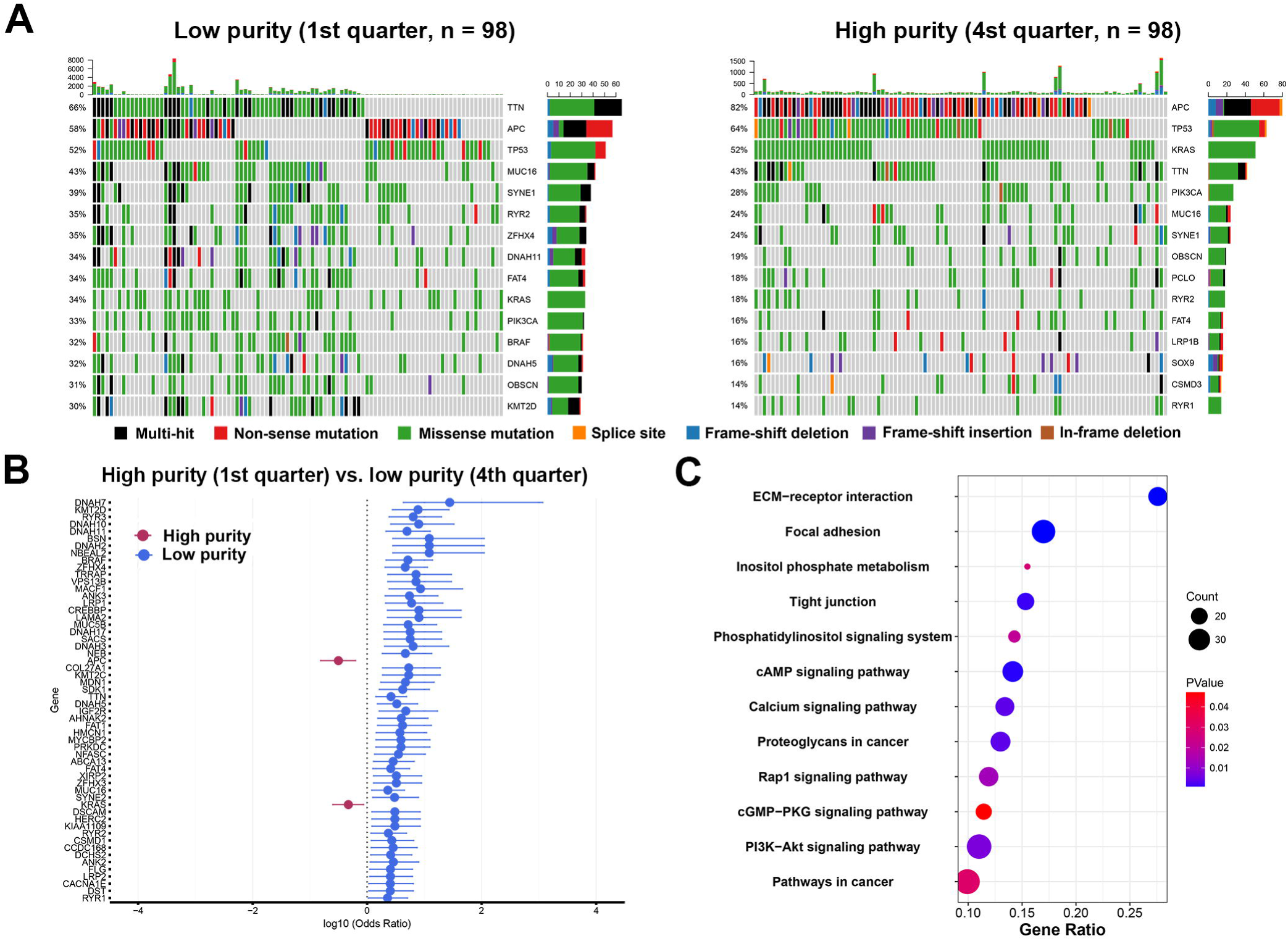
Mutation profile between low and high purity groups (1st vs. 4th quarter) in TCGA dataset. (**A**) Mutation profile of low and high purity groups. (**B**) Differentially mutated genes between low and high purity groups. (**C**) Highly mutated pathways in low purity group.

By performing KEGG analysis on significantly highly mutated genes in low purity groups, we further investigated pathways involving tumour purity. Pathways related to ECM-receptor interaction, focal adhesion, calcium signalling were significantly more mutated in low purity group in all parallel analyses (Fig. 4C, Figure S4C, and Figure S5C). Moreover, more mutations were enriched in PI3 K-Akt signalling pathway in CC patients with low tumour purity, which is a classic oncogenic pathway promoting CC progression (25).

**Fig. 4.**
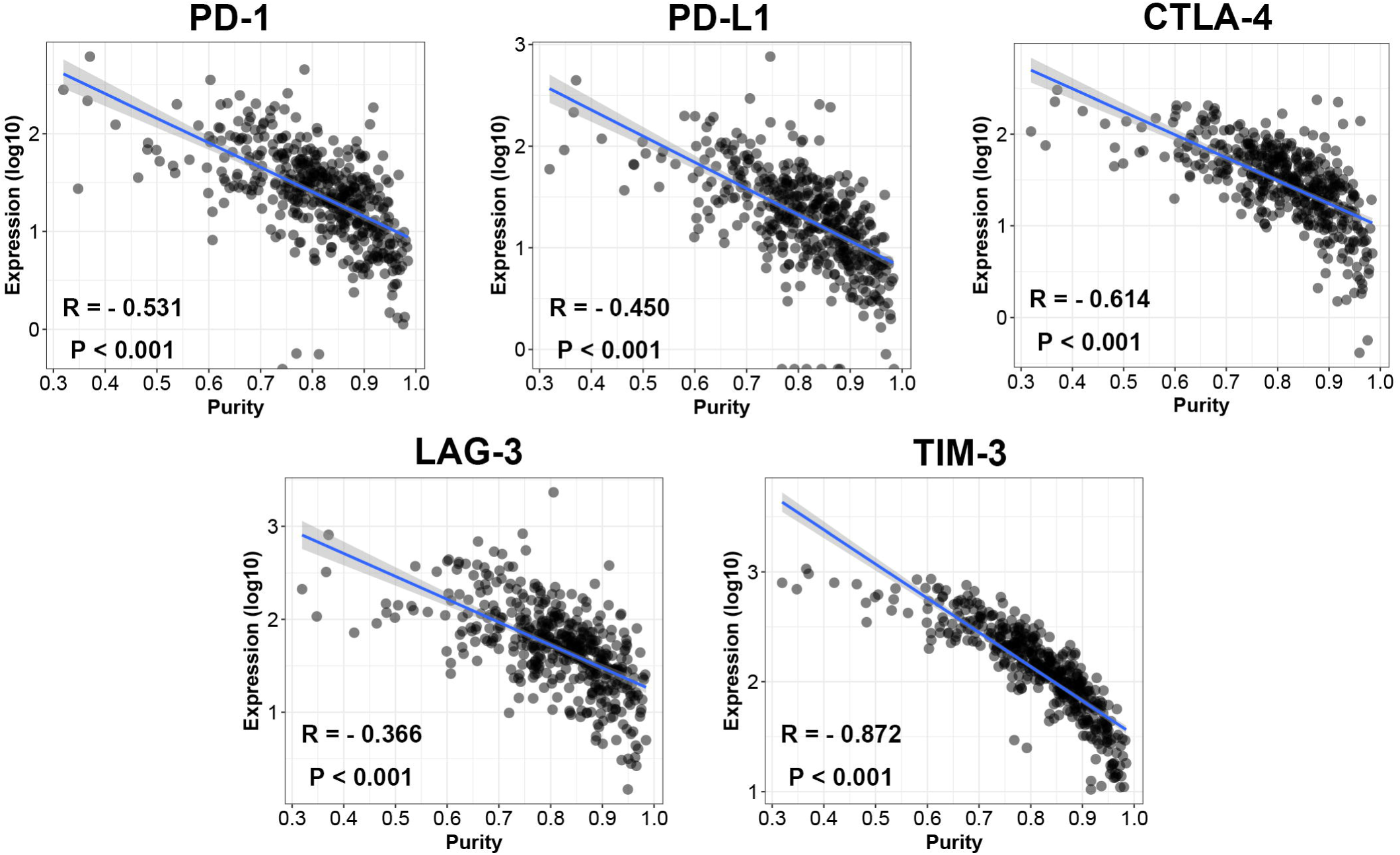
The expression level of immunotherapy-associated genes was inversely correlated with tumour purity in TCGA dataset.

The relationship between SCNAs and purity was also explored. SCNAs, including amplification, deletions and total SCNA events, were not associated with tumour purity (P = 0.547, P = 0.0648 and P = 0.0683, respectively) (Figure S6). The alteration pattern between low and high purity groups was similar (Figure S7). Parallel comparisons of SCNAs among subgroups (1st vs. 2nd half, 1st vs. 3rd tertile, 1st vs. 4th quarter) revealed no differential cytoband amplification or deletions using Fisher’s exact tests, which indicated that SCNAs might be irrelevant with CC purity.

### Tumour purity related immune microenvironment

To explore the underlying mechanism of purity’s prognostic value, we performed differential expression analysis on TCGA RNA-Seq data between low and high purity groups divided by median purity. Notably, the expression level of immunotherapy-associated markers (PD-1, PD-L1, CTLA-4, LAG-3, and TIM-3) was inversely correlated with tumour purity in all three datasets (Figure 4, Figure S8, and Figure S9), which was similar to a previous pan-cancer analysis (26). Genes with FDR < 0.05 in differential expression analysis were further accessed using GSEA. Multiple immune-related pathways were highly enriched in low purity group (Table S4), including positive regulation of inflammatory response, positive regulation of leukocyte mediated immunity, lymphocyte migration and adaptive immune response, etc. (Figure 5A) Therefore, low purity CC exhibited intensive immune phenotype.

**Fig. 5.**
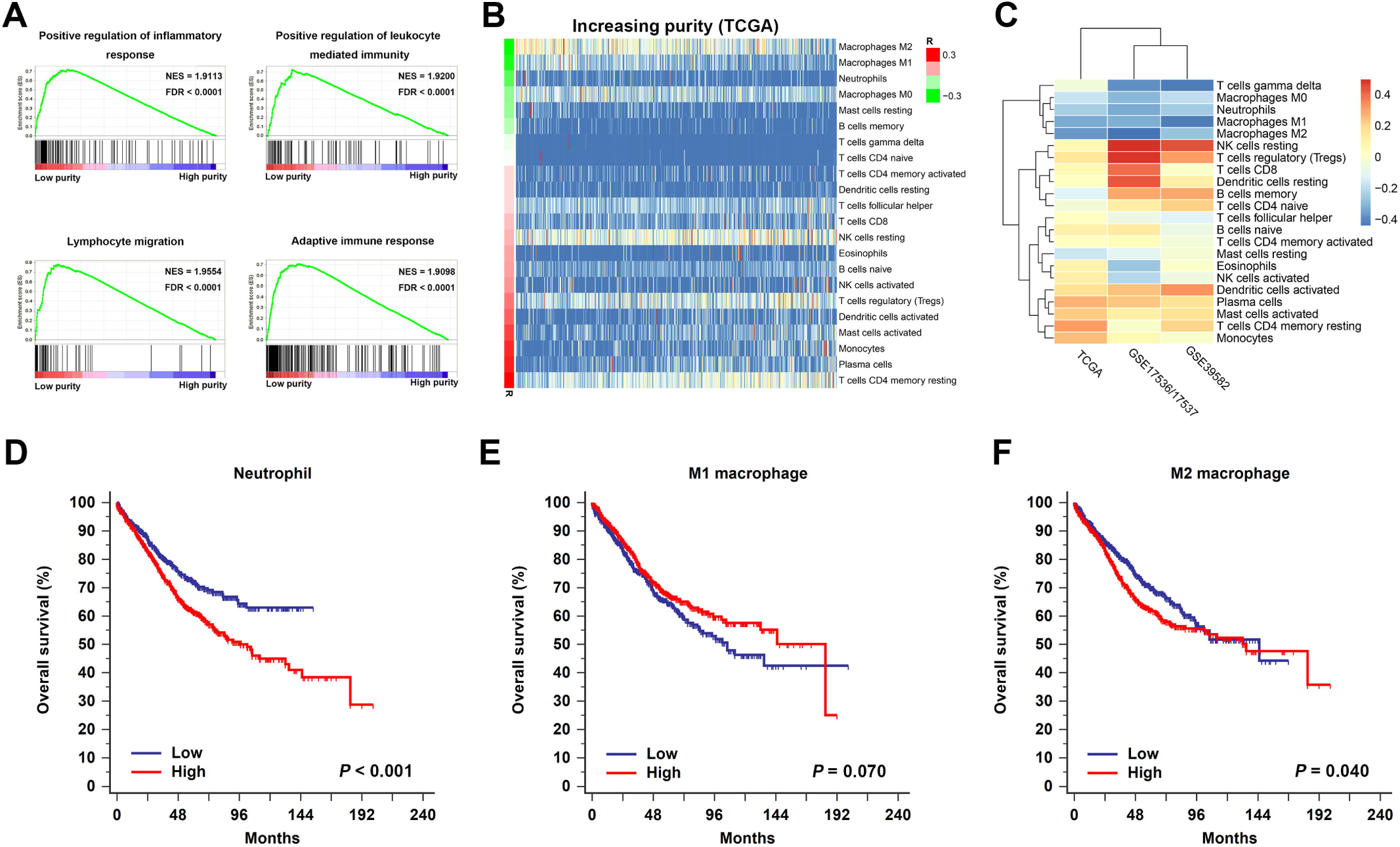
Microenvironment changing associated with tumour purity. (**A**) Immune-related pathways were highly enriched in low purity group. (**B**) The distribution of relative proportion of immune cells sorted by increasing purity in TCGA dataset. The left colour bar indicated the correlation coefficients. (**C**) The correlation coefficients of immune cells and tumour purity in TCGA, GSE17536/17537, and GSE 39582. (**D**) High relative proportion of neutrophils indicated poor overall survival. (**E**) The relative proportion of M1 macrophages was not significantly associated with overall survival. (**F**) High relative proportion of M1 macrophages predicted poor prognosis. NES: normalized enrichment score; FDR: false discovery rate.

As immune cells composed the major non-tumour proportion of microenvironment, we tried to figure out which types of tumour-infiltrating immune cells were related to tumour purity and immune phenotype. We applied CIBERSORT algorithm on RNA-Seq and microarray data to estimate the relative proportion of 22 immune cells of leukocytes for each CC sample. The association of different cell types and purity was plotted in Figure 5B, Figure S10, and Figure S11. Heatmap illustrating the correlation coefficients of immune cells and tumour purity was shown in Figure 5C. The absolute value of coefficients more than 0.2 across three datasets was applied as a threshold for filtration. Among 22 immune cell types, M1 macrophages, M2 macrophages, and neutrophils were negatively correlated with tumour purity in all three independent datasets (all R < −0.2). Next, samples from combined dataset were divided into low and high groups according to the corresponding median proportion of immune cells. By performing Kaplan–Meier survival analyses and Log-rank tests, the relative proportion of M1 macrophages was not significantly associated with OS (P = 0.070) (Figure 5E). However, the relative proportion of neutrophils and M2 macrophages were identified as indicators for poor prognosis (Figure 5D and Figure 5F) (*P* < 0.001 and P = 0.040, respectively), which may partially explain the worse prognosis in low purity CC.

## Discussion

In line with previous studies, we found low tumour purity, aka high stroma infiltrates, was negatively associated with shorter survival (6–10). However, we applied a computational method for purity speculation rather than histopathological way. In recent years, numerous computational tools for tumour purity estimation were proposed based on different types of genetic data (27–29). A comparative study showed a high concordance between methods (5). ESTIMATE algorithm was selected in our research for its compatibility of RNA-Seq data and microarray data.

Tumour stroma has been reported to facilitate tumour budding, epithelial-mesenchymal transition and lymphatic metastases (30,31). We found the tumour purity of stage III CC samples was relatively lower, implying the tumour purity may change dynamically during tumour progression. In addition, dMMR CC samples were found having significant lower tumour cell proportion than pMMR ones. dMMR CC is a special subtype of CC, which has increased mutational burden leading to neoepitopes production (32). Numerous aberrant proteins trigger an antitumor immune response. A close relationship of high immune cell infiltrate and dMMR has been demonstrated previously, which is consistent with our findings that low purity CC was characterized by the more dMMR percentage and increased mutation rate (33).

Previous histopathological based researches suggested that stroma percentage was negatively associated with adjuvant chemotherapy benefit (8,9). We further validated this finding by a computational approach. Subgroup analyses and Cox regression analyses indicated tumour purity as an independent prognostic factor in CC patients receiving adjuvant chemotherapy. Researches targeting TME developed fast recently, especially checkpoint blockade immunotherapy which shows therapeutic potential in dMMR CRC patients (34,35). Interestingly, immunotherapy-associated markers (PD-1, PD-L1, CTLA-4, LAG-3, and TIM-3) were high expressed in low purity CC. Therefore, addition of TME-targeted therapy to current 5-FU based adjuvant chemotherapy may be an option to improve oncologic outcomes for low purity population (36).

Gene level association of tumour purity was also interpreted in this study. The mutation burden in low purity CC was significantly heavier than high purity ones, which was consistent with the previous reports in pan-cancer analysis and glioma (5,16). Notably, Only *APC* and *KRAS* mutations showed increased frequency in high purity CC. KEGG analysis revealed that pathways related to extracellular matrix (ECM) were highly mutated in low purity CC samples. The abnormality of ECM contributes to cancer progression, invasion, and metastasis (37). Mutations were also differentially enriched in the PI3 K-Akt pathway in low purity group, which is a crucial pathway in CC progression and chemotherapy resistance (38). The increasing mutations in above pathways may contribute to the impaired survival in low purity CC patients.

Frequent SCNA events were observed in CC, which may lead to gene expression alterations and promote cancer progression (14,39). Unlike glioma, where increased SCNA events were detected in high purity tumours, SCNAs in CC were not significantly associated with purity (16). In line with our findings, a recent report showed that for microsatellite stable (MSS) CC, stroma proportion was not correlated with SCNA events (40). As high stromal proportion may disturb the measurement of SCNAs, the correlation between tumour purity and SCNAs need to be further investigated.

GSEA analyses revealed that immune-related pathways were highly enriched in low purity tumours. Furthermore, tumour purity was found negatively correlated to the relative proportion of neutrophils, M1 macrophages and M2 macrophages in all three independent datasets. Among three types of immune cells, the relative proportion of neutrophils and M2 macrophages presented negative prognostic value, which may explain the worse survival in low purity CC. The adverse association between neutrophils and M2 macrophages infiltration and CC prognosis has long been noticed. Unlike the protumoural effect of M2 macrophages was accepted as a general idea, the role of neutrophil infiltration in CRC prognosis remains uncertain, both beneficial and harmful effect of neutrophils was observed in previous studies (41–46). Heterogeneous markers including CD66b, CD177 or myeloperoxidase were used for neutrophil identification, which may bias the conclusions (43–46). CIBERSORT, which utilizes clusters of genes for cell separation from RNA mixtures, may be more suitable for immune cell quantification than traditional IHC staining of limited markers. Several limitations of our study need to be noticed. Purity estimation in our investigation was only calculated by one computational method based on transcriptomic data, further validation using multiple algorithms based on mutation, CNA or DNA methylation data may be needed. Moreover, due to the retrospective setting of this study, prospective researches are required to further access our conclusions.

In summary, we systematically evaluated the role of tumour purity in CC prognosis, gene profile and microenvironment. Tumour purity was identified as an adverse independent prognostic factor in CC, as well as an independent predictor of adjuvant chemotherapy benefit. Low purity CC exhibited heavier mutation burden but with lower APC and KRAS mutation frequency. Nevertheless, SCNA was not significantly correlated with tumour purity. Immune-related pathways were highly enriched in low purity group. Furthermore, M2 macrophages and neutrophils, which indicated worse survival in CC, were negatively associated with purity. The role of tumour purity in CC needs further investigation for better prognostic stratification and clinical management.

## Supplementary material

Supplementary data are available at http://carcin.oxfordjournals.org/

